# The geometry of dominance shows broad potential for stable polymorphism under antagonistic pleiotropy

**DOI:** 10.64898/2026.03.27.714876

**Authors:** Evgeny Brud, Rafael F. Guerrero

## Abstract

Alleles with opposing effects on fitness characters are said to exhibit selectional antagonistic pleiotropy (broadly construed so that effects are not necessarily confined to the same individual). A number of theoretical investigations considered the case where a pair of alleles at a locus influences two fitness components and derived the conditions giving rise to stable polymorphism under various assumptions about the mode of trait-interaction. Strikingly, many of these analyses concluded that the potential for maintaining polymorphism is strongly constrained by the joint influence of two factors: (1) the prevalence of weak selection coefficients over coefficients of large magnitude, and (2) the absence of beneficial dominance reversals (where the deleterious effects of each allele are partially or completely masked in the heterozygous genotype). Consequently, the conclusion that selective polymorphism is unlikely to be maintained by intralocus mechanisms of antagonistic pleiotropy has achieved widespread acceptance. Here we argue that such conclusions do not apply to any of the following models of antagonism: (i) additive trait-interaction, (ii) multiplicative trait-interaction, (iii) bivoltine selection, (iv) soft selection, (v) hard selection, and (vi) sexual antagonism. We demonstrate that the parameter space giving rise to stable allelic variation is quite large throughout, and moreover, the plenitude of suitable parameters neither depends on the strength of selection nor requires dominance reversal. Dominance coefficients associated with stringent conditions for stable polymorphism are shown to be atypical as compared to all feasible parameters, and best regarded as an outcome of adherence to a special relation: dominance with a constant magnitude and direction, which includes the case of additive allelic effects at a locus. Properties of single-locus equilibria (heterozygosity, allele frequency differentiation) are investigated, as well as the contribution of dominance schemes to the genetic variance in fitness characters in populations at multilocus linkage equilibrium.

**Author summary:** Allelic variants at a locus with opposing effects on multiple fitness components (antagonistic fitness pleiotropy) have long been appreciated as a possible source of balancing selection. The prevalence of polymorphism owing to this form of natural selection, however, has been doubted on theoretical grounds due to the fact that standard assumptions of genetic models (namely, constant magnitudes for the dominance coefficients) are hardly conducive to the maintenance of polymorphism. The major exception to this conclusion lies with schemes that exhibit dominance reversal (where the direction of dominance for antagonistic alleles flips across fitness components). Here we conduct a geometric analysis of the space of polymorphism-promoting dominance parameters and conclude that the conditions for maintaining balanced alleles is unrestrictive, with non-reversals playing an underappreciated role.

## Introduction

A gene with effects on more than one phenotypic character is said to exhibit *pleiotropy*; effects are defined as concordant when codirectional and antagonistic when in opposition. Examples of pleiotropy are widespread (see [1] for a recent review), and antagonistic effects feature prominently in evolutionary explanations for diverse phenomena including aging [2, 3], sexual dimorphism [4, 5], and genome evolution (e.g., sex chromosome differentiation; 6, 7, 8). Antagonistic selection has also received significant attention in population genetics theory as a mechanism for maintaining balanced polymorphism [9, 10, 11, 12, 13, 14, 15]. Despite this broad theoretical and conceptual importance, it remains unclear whether antagonistic fitness pleiotropy substantially contributes to the maintenance of variation in natural populations.

To address this question, classic theoretical work explored the simplest diploid genetic architecture for antagonistic pleiotropy (AP): a single locus with two alleles. Deterministic analyses typically derived conditions for protected polymorphism [16, 17], i.e., conditions under which *Aa* heterozygotes can invade either class of homozygous resident population (*AA* or *aa*). Elegant expressions for the suitable space of polymorphism-promoting parameters were thereby derived for a number of diverse scenarios of antagonistic selection, of which we focus on the following six: (1) additive fitness pleiotropy [9, 10], (2) multiplicative fitness pleiotropy [9, 10], (3) seasonal selection in a bivoltine population (i.e. organisms with life cycles producing two generations per year; [18, 19, 20, 21, 22]), the multiple-niche models of (4) soft selection and (5) hard selection [16, 23, 24, 25], and (6) sexual antagonism [26, 27].

The implications of these theoretical analyses were themselves somewhat antagonistic in character. On the one hand, these models were conceptually groundbreaking in demonstrating that the maintenance of polymorphism extends beyond the canonical case of single-locus heterozygote advantage, in which heterozygotes are directly fitter than either homozygote. They showed that stable polymorphism can emerge from the interaction of multiple fitness components, allowing antagonistic pleiotropy to balance alleles even when no single component exhibits overdominance and under conventional dominance schemes spanning partial dominance through recessivity.

On the other hand, severe skepticism soon emerged because these models appeared to permit stable polymorphism only under restrictive parameter combinations, raising doubts about their relevance in natural populations. Hedrick [28] argued that under constant dominance schemes, in which the dominance relation between alleles remains the same across antagonistic fitness components (also called parallel dominance in Curtsinger [10] and cumulative overdominance in Bertram et al. [21], after Dempster [18]), the region of selection coefficients that supports polymorphism is rather tight. In fact, across all six focal models, constant dominance and additivity (the schemes most commonly assumed in theoretical work) confine polymorphism to cases where selection coefficients (*s*_1_, *s*_2_) are nearly symmetric (*s*_1_ ≈ *s*_2_) in the limit of weak selection, with departures from symmetry arising only when fitness differences are unusually large (∼ 20% selection coefficients or greater). As summarized by [28, p. 131], “[i]n general, only when the selective differences are large and fairly similar in size is the likelihood of a stable polymorphism from antagonistic pleiotropy significant. The only major exception to these conclusions appears to be when there is a reversal of dominance such that the heterozygote is similar in fitness to the favourable homozygote for both traits.” These sentiments were broadly shared [e.g., 9, 10], and prospects for the maintenance of polymorphism shifted to focusing on mechanisms by which reversing dominance schemes could underlie balancing selection (e.g., cyclical selection [12, 21, 18]; sexual antagonism [27]). These tended to highlight the special relation of symmetric dominance reversal, in which the alleles switch the direction of dominance across the two fitness components while maintaining similar magnitude. The limiting case of complete beneficial reversal, in which heterozygous fitness components are equal to that of the fittest homozygous component, was recognized as the optimal dominance scheme and therefore an especially potent stabilizer of allelic variation under general selection.The stability-conferring properties of reversing dominance were counterposed with the view that this genetic scheme, in the main, lacked a solid evidentiary basis to serve as a plausible and widespread mechanism for maintaining allelic variation [10]; cf. [29], particularly when dominance relations were construed as being strongly determined by biochemical and physiological principles such that dominance coefficients between allelic pairs were expected to be rigid and fixed, as opposed to pliant and modifiable [30, 31]; cf. [32].

Here we argue that the view of antagonistic pleiotropy (AP) as a highly restrictive mechanism for maintaining polymorphism is unwarranted, and that the resulting emphasis on dominance reversal as a stabilizing force is correspondingly misplaced. By examining the full space of feasible fitness parameters without imposing special relations on allelic dominance (i.e., constant dominance, additivity, or symmetric dominance reversal), we demonstrate that the full parameter space is broadly conducive to the maintenance of polymorphism in all six models, with only minor quantitative differences between them. The probability of encountering stabilizing parameter combinations remains substantial across a broad range of selection strengths, including fairly weak selection, indicating that the “tight interval effect” described above is not a general feature of antagonistic pleiotropy. Moreover, non-reversing dominance contributes substantially throughout, implying that no *a priori* emphasis need be placed on dominance reversals as the predominant mechanism balancing antagonistic alleles. A comparison of allele frequencies at equilibrium shows substantial heterozygosity is maintained even when antagonistic selection coefficients differ by up to two-fold in magnitude. Under weak selection, the stabilizing regions of parameter space converge across all six models, forming a common domain conducive to polymorphism, whereas stronger selection enlarges this domain only modestly, with its primary effect expressed through increased allele-frequency differentiation. Lastly, we derive the additive and dominance genetic variances for equilibrium states in which *L* loci segregate independently (multilocus linkage equilibrium) and show that both non-reversing and reversing dominance stabilize substantial levels of additive variance in fitness components.

## Models

### Single-locus models

The diploid models of antagonistic pleiotropy (AP) assume (i) there exist two fitness characters under the control of a single biallelic autosomal locus (Table 1), (ii) mating occurs at random, and (iii) genetic drift is absent (see Fig. S1 for life-cycle diagrams). Deterministic equations (summarized in Appendix A) govern the change in allele frequencies between successive generations for *A* and *a* (initially *p* and *q*, respectively, with *p* + *q* = 1) for the generalized fitness parameters in Table 1).

**Table 1.** Fitness component values for genotypes subject to two-trait antagonistic selection.

| Parameters | Component | AA | Aa | aa |
| --- | --- | --- | --- | --- |
| General | 1 | $W_{AA,1}$ | $W_{Aa,1}$ | $W_{aa,1}$ |
| | 2 | $W_{AA,2}$ | $W_{Aa,2}$ | $W_{aa,2}$ |
| Bounded | 1 | 1 | $1 - h_1 s_1$ | $1 - s_1$ |
| | 2 | $1 - s_2$ | $1 - h_2 s_2$ | 1 |

$W_{\text{genotype},i} \in \mathbb{R}_+$ , $s_1, s_2 \in (0, 1)$ , and $h_1, h_2 \in [0, 1]$ .

### Conditions for protected polymorphism

Protected polymorphism requires that the mutant allele increase when rare but decline when nearly fixed. In other words, if the relative fitness of the *a*-allele is 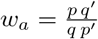 sufficient conditions for protection are lim_*q*→0_ *w*_*a*_ > 1 and lim_*q*→1_ *w*_*a*_ < 1. Additional protection conditions exist when one of the invasion fitnesses is neutral and the derivative of invasion fitness at the boundary is positive [17]. Because these cases require exact equality among parameter values rather than defining regions of parameter space with positive area, we do not consider them further.

Sexual antagonism is the only one of our focal models with 2 variables being tracked at the census points of the life cycle. Thus, the single-variable criterion (positive allele frequency change between the initial and second generation) no longer suffices because there is the logical possibility that a non-positive allele frequency change in the initial two generations coincides with changes in the genotypic configurations (i.e. deviations from HWE). Consequently, the *a*-allele might have a long-run advantage in some later genotypic environments as opposed to the initial two generations. This possibility does not arise in the other models. Thus, under sexual antagonism the invasion fitness of rare mutants equals the magnitude of the dominant eigenvalue of the following Jacobian matrices evaluated at the boundary equilibria:

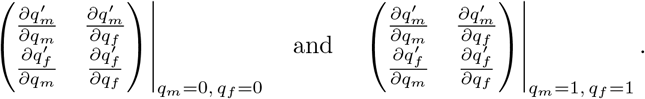

These turn out to be equal to 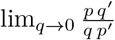 and 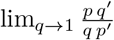 for the monomorphic states *q* = 0 and *q* = 1, respectively.

### Geometry of polymorphic space

The region of parameter space giving rise to protected polymorphism at a biallelic locus can be expressed in either of two informative formulations, namely in terms of the generalized fitness component parameters *W*_genotype,*i*_ or in terms of the selection and dominance coefficients (*s*_1_, *s*_2_, *h*_1_, *h*_2_). Assuming the fitness component values are positive and unbounded, *R*_*W*_ gives the generalized protection region:

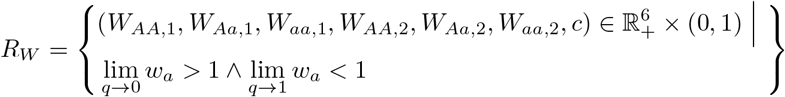

where the additional parameter bound that applies under two-niche selection, *c*, is the fraction of the total population that resides in subpopulation 1 prior to viability selection (*c* is vacuous in Models 1–3 and 6).

As cataloged in Table 2, a heterozygote advantage in the sum of fitness components defines the protection conditions under additive AP (Model 1), while a heterozygote advantage in their product determines these conditions under both multiplicative AP (Model 2) and bivoltine selection (Model 3) [9, 10, 28, 19, 21, 22]. The criteria for the soft selection (Model 4) and hard selection (Model 5) scenarios of two-niche antagonism involve weighted fitness calculations as well as ratios of fitness components (i.e., heterozygous fitness components relative to one or the other homozygote). Under soft selection, the sum of the size-weighted ratios of fitness components must exceed 1 for each homozygote comparison (Table 2; [16]). By contrast, under hard selection (where gamete-pool contributions from each niche are proportional to mean subpopulation fitness, rather than held constant) the focal comparison is the ratio of the size-weighted sums of fitness components (Table 2; [25]). The protection conditions for sexual antagonism (Model 6) amount to a special case (*c*_*i*_ = 1*/*2) of the soft selection condition, that is, with niches of equal size; this special case also results in an equivalence between the additive AP and hard selection conditions.

**Table 2.** Conditions for protected polymorphism at loci under antagonism.

| Model | Polymorphism conditions<br>($\lim_{q \rightarrow 0} w_a > 1 \wedge \lim_{q \rightarrow 1} w_a < 1$) |
| --- | --- |
| 1) Additive AP | $\sum_{i=1}^2 W_{Aa,i} > \sum_{i=1}^2 W_{aa,i} \wedge \sum_{i=1}^2 W_{Aa,i} > \sum_{i=1}^2 W_{AA,i}$ |
| 2) Multiplicative AP |  |
| *and* |  |
| 3) Bivoltine selection | $\prod_{i=1}^2 W_{Aa,i} > \prod_{i=1}^2 W_{aa,i} \wedge \prod_{i=1}^2 W_{Aa,i} > \prod_{i=1}^2 W_{AA,i}$ |
| 4) Soft selection* | $\sum_{i=1}^2 c_i \frac{W_{Aa,i}}{W_{AA,i}} > 1 \wedge \sum_{i=1}^2 c_i \frac{W_{Aa,i}}{W_{aa,i}} > 1$ |
| 5) Hard selection* | $\frac{\sum_{i=1}^2 c_i W_{Aa,i}}{\sum_{i=1}^2 c_i W_{AA,i}} > 1 \wedge \frac{\sum_{i=1}^2 c_i W_{Aa,i}}{\sum_{i=1}^2 c_i W_{aa,i}} > 1$ |
| 6) Sexual antagonism | $\frac{1}{2} \left( \sum_{i=1}^2 \frac{W_{Aa,i}}{W_{AA,i}} \right) > 1 \wedge \frac{1}{2} \left( \sum_{i=1}^2 \frac{W_{Aa,i}}{W_{aa,i}} \right) > 1$ |

\*Where $c_1 = 1 - c_2 = c$ .

With *W*_*AA*,1_ = 1 and *W*_*aa*,2_ = 1 (and substituting the values from Table 1), *R*_*W*_ can be reformulated to give the bounded region *R*_*hs*_, to the exclusion of any scenario involving overdominance, underdominance, neutrality, or lethality with regard to any single fitness component. This region is

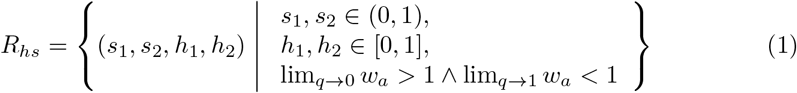

(for some *c*, in Models 4–5). Without loss of generality, we assume *s*_1_ ≤ *s*_2_. Using the `Reduce` function in *Mathematica* (version 14.3; Wolfram [33]), *R*_*hs*_ is equal to

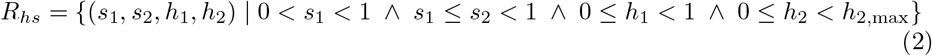

(for some *c*, in Models 4–5), where the upper boundary on component-2 dominance, *h*_2,max_, is a function of (*s*_1_, *s*_2_, *h*_1_) and is always less than or equal to 1 (Table 3). Given any (*s*_1_, *s*_2_) and assuming *c* = 1*/*2, the conditions for protected polymorphism reduce to conditions on dominance that define the region *P*:

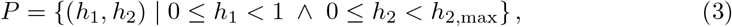

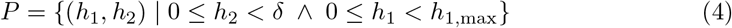

where 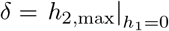, and *h*_1,max_ is a function of (*s*_1_, *s*_2_, *h*_2_) and is always less than or equal to 1 (Table 3).

**Table 3.** Upper boundaries on dominance coefficients in polymorphic region *P*.

| Model | $h_{1,\max}$ | $h_{2,\max}$ |
| --- | --- | --- |
| 1) Additive AP |  |  |
| and | $\frac{s_1 - h_2 s_2}{s_1}$ | $\frac{s_1 - h_1 s_1}{s_2}$ |
| 5) Hard selection with $c = \frac{1}{2}$ | | |
| 2) Multiplicative AP |  |  |
| and | $\frac{s_1 - h_2 s_2}{s_1 - h_2 s_1 s_2}$ | $\frac{s_1 - h_1 s_1}{s_2 - h_1 s_1 s_2}$ |
| 3) Bivoltine selection |  |  |
| 4) Soft selection with $c = \frac{1}{2}$ | $\begin{cases} \frac{(1 - h_2)s_2}{s_1(1 - s_2)} & \text{if } H_2 \\ 1 - \frac{h_2 s_2(1 - s_1)}{s_1} & \text{otherwise} \end{cases}$ | $\begin{cases} 1 - \frac{h_1 s_1(1 - s_2)}{s_2} & \text{if } H_1 \\ \frac{(1 - h_1)s_1}{(1 - s_1)s_2} & \text{otherwise} \end{cases}$ |
| and |  |  |
| 6) Sexual antagonism |  |  |

where $H_1 : s_1 \leq s_2 < \frac{s_1}{1-s_1} \wedge h_1 \leq \frac{2}{s_1+s_2-s_1s_2} - \frac{1}{s_1}$ , and $H_2 : s_1 \leq s_2 < \frac{s_1}{1-s_1} \wedge \frac{s_1-(s_1+1)s_2}{(s_1(s_2-1)-s_2)s_2} < h_2$ .

The set *P* delimits a geometric region on the unit square of dominance coefficients with vertices at (0, 0), (1, 0), and (0, *δ*), where the origin (i.e. complete beneficial reversal; always an element of *P*) is connected to the points (1, 0) and (0, *δ*) (not elements of *P*) by line segments. Moreover, (1, 0) and (0, *δ*) are connected by the upper dominance boundary (see the alternative expressions in each row of Table 3). In all cases, these upper boundaries are strictly decreasing functions of their permissible dominance domains and are (i) affine under additive protection conditions (Models 1,5), (ii) concave under multiplicative conditions (Models 2,3), and (iii) concave as well as piecewise affine for the models featuring weighted harmonic mean conditions on homozygous fitness relative to that of heterozygotes (Models 4,6) (Fig. 1).

**Fig 1.**
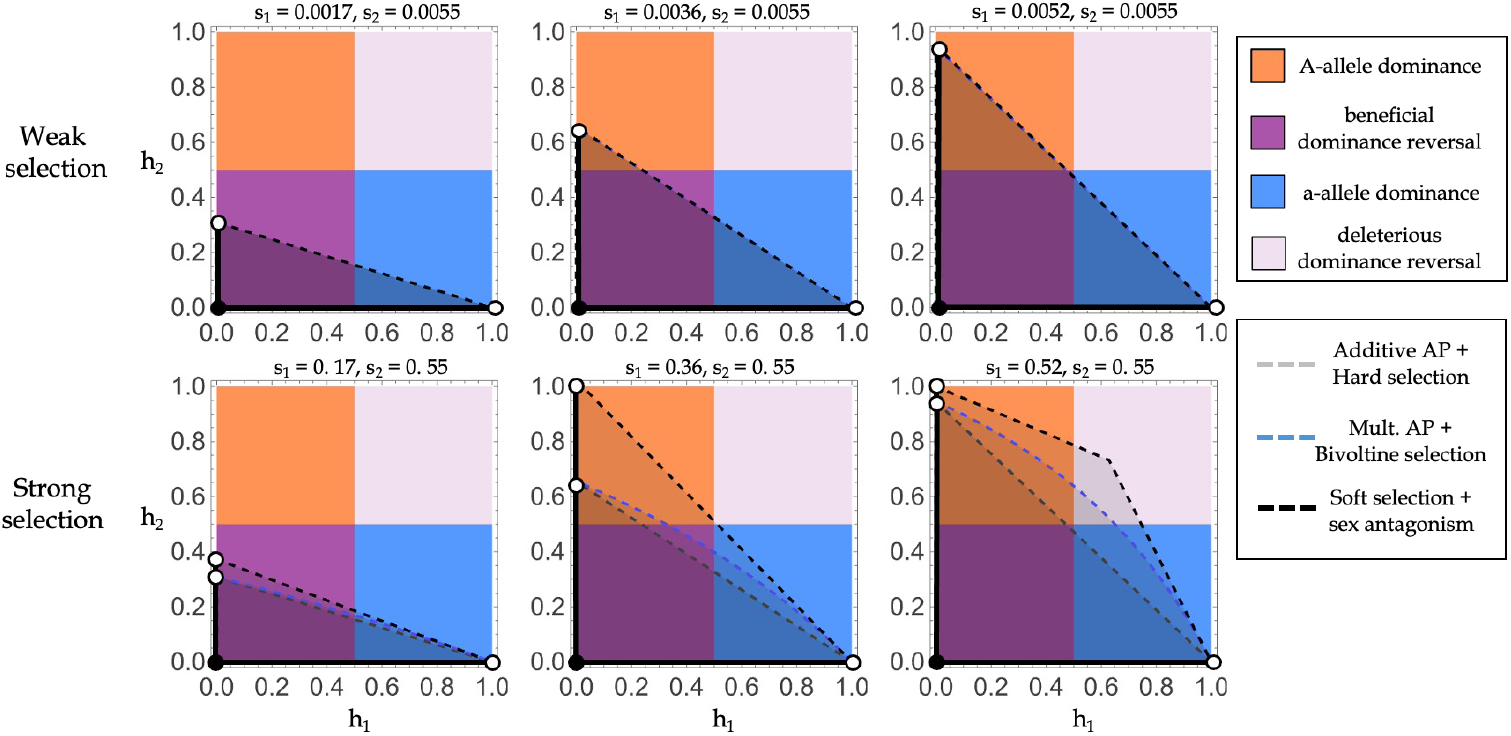
Polymorphism-promoting dominance schemes form the region *P* on the unit square of dominance coefficients. Each quadrant of the unit square corresponds to a unique category of component-specific dominance (top legend). The region *P* (shaded portion) confers protected polymorphism for the pair of selection coefficients indicated on the plot labels; bold lines and points denote inclusion in *P* while dashed lines and open circles denote exclusion. The dashed boundaries (bottom legend) correspond to the model-specific upper dominance boundaries listed in Table 3. **Top row:** The intercept on the *h*_2_-axis equals the ratio *s*_1_*/s*_2_. **Bottom row:** Strong selection coefficients result in considerable expansion of the *P* region, especially for selective regimes with less than two-fold asymmetry in *s*_1_, *s*_2_ and non-additive component interaction. The intercept on the *h*_2_-axis equals the ratio *s*_1_*/s*_2_ under all models except for soft selection and sexual antagonism, for which the intercept is the larger value.

The size of *P* is a measure of the potential for biallelic polymorphism and is equivalent to the probability that random uniform sampling of dominance parameters from the unit square results in a protected polymorphism [34, 35]. Moreover, the distribution of *P* across the quadrants of the unit square provides the proportional contributions of the four major categories of two-component dominance (A-allele dominance, *a*-allele dominance, beneficial dominance reversal, and deleterious dominance reversal) to the overall potential. The area formulas for *P* are provided in Table 4 and the formulas for the quadrant-specific intersections of *P* are provided in Appendix B.

**Table 4.** Potential for biallelic polymorphism.

| Model | $\text{Area}(P) = \int_0^1 h_{2,\max} dh_1 =$ |
| --- | --- |
| 1) Additive AP |  |
| *and* | $\frac{s_1}{2s_2}$ |
| 6) Hard selection with $c = \frac{1}{2}$ | |
| 2) Multiplicative AP |  |
| *and* | $\frac{s_1 + (1 - s_1) \ln(1 - s_1)}{s_1 s_2}$ |
| 3) Bivoltine selection |  |
| 4) Soft selection with $c = \frac{1}{2}$ | $\begin{cases} \frac{s_1}{2(1 - s_1)s_2} & \text{if } \Omega_1 \\ \frac{s_2^2 - (s_2 - 1)s_1^2 - s_2(s_2 + 2)s_1}{2s_1(s_1(s_2 - 1) - s_2)s_2} & \text{if } \Omega_2 \end{cases}$ |
| *and* |  |
| 5) Sexual antagonism |  |

where $\Omega_1 : 0 < s_1 < \frac{1}{2} \wedge \frac{s_1}{1-s_1} < s_2 < 1$ , and $\Omega_2 : 0 < s_1 < 1 \wedge s_1 \leq s_2 < \frac{s_1}{1-s_1}$ .

### Equilibrium allele frequencies and differentiation

The solution of *w*_*a*_ = 1 in terms of the single variable *q* (over the reals on the unit interval) provides the stable equilibrium frequency of the *a*-allele, 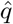. The only exception to this is Model 6 (sexual antagonism), where the equations involve two variables (*q*_*m*_, *q*_*f*_) and the system to solve is 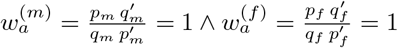.

For Models 3–6, the equilibrium state can always be decomposed into two “subgroup” allele frequencies, 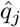 and 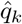 (i.e. where 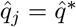 and 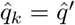 are the season-specific frequencies under bivoltine selection (Model 3); 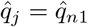 and 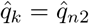 are the niche-specific (post-selection) frequencies under spatial selection (Models 4–5); and 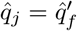 and 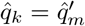 provide the (post-selection) sex-specific frequencies under sexual antagonism (Model 6)).

Moreover, for each model 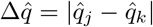 provides an absolute measure of allele-frequency differentiation between subgroups. Models 1–2 lack a relevant basis for differentiating subgroups, and so trivially 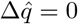 in these scenarios.

### Multilocus extension under linkage equilibrium

We start from the additive and dominance genetic variances (at a single locus) for each of two fitness characters provided by Rose [9], in which the parameterization for the genotypic fitness effects inn a component are given as deviations from the heterozygous value (Table 5). The additive genetic variances are *V*_*A*1_ = 2*pq ε*^2^(*η*_1_*p* + *q*)^2^ and *V*_*A*2_ = 2*pq δ*^2^(*p* + *η*_2_*q*)^2^ for fitness component 1 and 2, respectively. The corresponding dominance variances are *V*_*D*1_ = *p*^2^*q*^2^*ε*^2^(1 − *η*_1_)^2^ and *V*_*D*2_ = *p*^2^*q*^2^*δ*^2^(1 − *η*_2_)^2^, where *η*_*i*_ is the dominance parameter for component *i*; note that *η*_*i*_ ∈ [0, ∞), and additivity corresponds to *η*_*i*_ = 1. The parameters *ε* and *δ* denote the additive fitness effects on components 1 and 2, respectively. A translation between Rose’s (1982) parameterization and the bounded formulation of Table 1 is obtained by first relativizing the fitnesses to the corresponding (row-specific) heterozygous genotype.

**Table 5.**
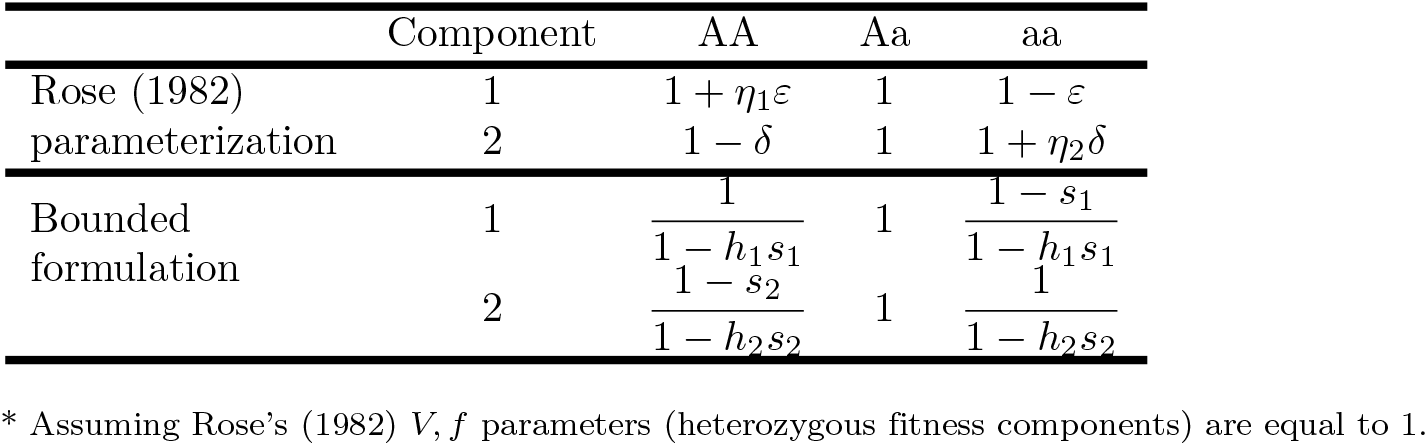
Fitness component values under two-trait antagonistic selection.

Equating the corresponding entries of the parameterization schemes in Table 5 provides the identities 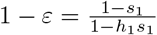 and 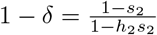, so the additive effects are 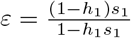 and 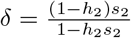. Via substitution of *ε* and *δ*, we can solve for the dominance parameters 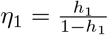, and 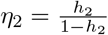.

Assuming the alleles have already settled to their single-locus equilibrium values 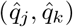, we can reformulate the additive and dominance genetic variances in terms of the selection coefficients and dominance coefficients of Table 1:

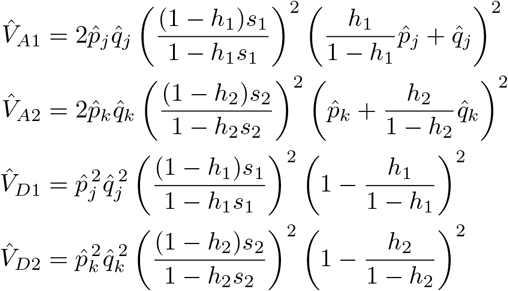

(Note that the equilibrium frequencies are also in terms of *h*_1_, *s*_1_, *h*_2_, *s*_2_ upon evaluation; see Results.) The total genetic variances are 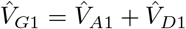 and 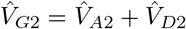. In a panmictic population subject to antagonistic selection at *L* loci, in which linkage equilibrium is constant and alleles at different loci act on fitness independently, the single-locus expressions above can be indexed by locus *i* and summed to provide the multilocus additive and dominance genetic variances at equilibrium (e.g. 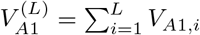).

### Additive and dominance variance at multilocus equilibrium

It is a priori unclear whether reversing and non-reversing dominance differ in their contribution to the levels of additive genetic variance in fitness components at equilibrium. In the terminology of quantitative genetics, these categories entail different sets of dominance deviations (departures of additive intralocus effects) as well as different levels of equilibrium allele frequencies (as demonstrated in the previous section), and it is these quantities, together with the additive effects, that determine the standing levels of genetic variance in fitness characters at equilibrium.

Rose (1982) provided formulas for the additive 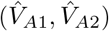 and dominance 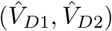 genetic variances for components 1 and 2, but did not evaluate them at equilibrium despite having also derived expressions for equilibrium allele frequencies under additive AP and multiplicative AP.

Recall that the single-locus equilibria under all six models converge to the expectation for additive AP in the limit of small selection coefficients (Fig. S3, Appendix C), and furthermore the allele-frequency differentiation converges to zero in the weak-selection limit (since *s*_1_, *s*_2_ determine the strength of differentiation).

Translating Rose’s (1982) parameterization into that of Table 5 and evaluating at the additive AP equilibrium

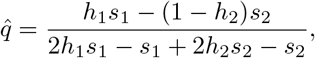

the approximate additive genetic variances for all six models under weak selection are

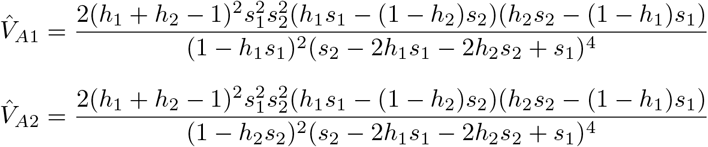

The approximate dominance genetic variances are

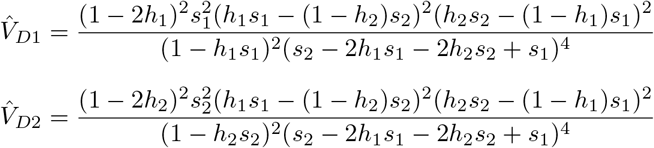

Given that additive intralocus effects (*h*_1_ = *h*_2_ = 0.5) are not conducive to the maintenance of single-locus polymorphism (i.e. this dominance scheme is generally excluded from the *P* region under weak selection), we investigate the effect of dominance categories on genetic variance by taking complete beneficial dominance reversal as the reference for comparison.

Evaluating these terms at *h*_1_ = *h*_2_ = 0 gives

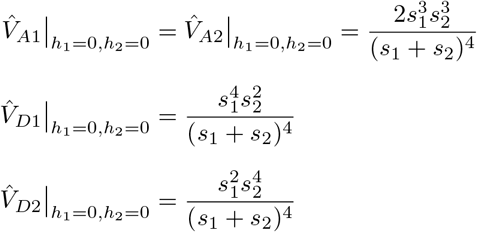

## Results

We re-derived the parameter regions conferring stable polymorphism by formulating the genotype-specific fitness components in terms of selection coefficients(*s*_1_, *s*_2_), and dominance coefficients (*h*_1_, *h*_2_). These conditions reduce to a characteristic region *P* on the unit square of dominance coefficients (Fig. 1). The *P* region is exactly identical between certain models: Additive AP and Hard selection (*c* = 1*/*2); Multiplicative AP and Bivoltine selection; Sexual antagonism and Soft selection (*c* = 1*/*2).

In fact, for “weak” selection (construed rather liberally so that *s*_1_, *s*_2_ ∼ 0–1%), *P* results in nearly complete overlap for all six models and takes the form of a right triangle with its right angle at the origin and with vertices at (0, *δ*) and (1, 0), where *δ* ≈ *s*_1_*/s*_2_ (Fig. 1, top row; Appendix C). Very strong selection regimes result in an expansion of this region beyond the right-triangle shape by intensifying the concavity of the upper dominance boundary (the “hypotenuse” present under weak selection), except in the cases of Additive AP and Hard selection (*c* = 1*/*2), whose protection regions retain the right-triangle shape throughout (Fig. 1, bottom row).

The size of this parameter region and its distribution over the quadrants of the dominance unit square provide, respectively, the potential for polymorphism of two alleles at a locus as well as the relative contribution of dominance categories (beneficial reversal, *A*-allele dominance, *a*-allele dominance, and deleterious reversal) to the overall potential (Appendix 2). For polymorphisms in *P*, we also investigate properties of the equilibrium state, including allele frequency, allele-frequency differentiation between focal subgroups of the population, as well as the levels of additive genetic variance in fitness components.

### Potential for polymorphism

The potential for polymorphism is substantial for all but the most asymmetric pairs of (*s*_1_, *s*_2_) in all six models. There is at least a 10% probability of randomly sampling a stabilizing dominance scheme for ∼ 80% of the possible pairs of (*s*_1_, *s*_2_); the probability increases from 10% to 50% with greater symmetry of the selection coefficients (Fig. 2A). Under strong selection, the polymorphic potential can exceed 50% for all models except Additive AP and Hard selection (these models are entirely insensitive to the strength of selection in this regard).

**Fig 2.**
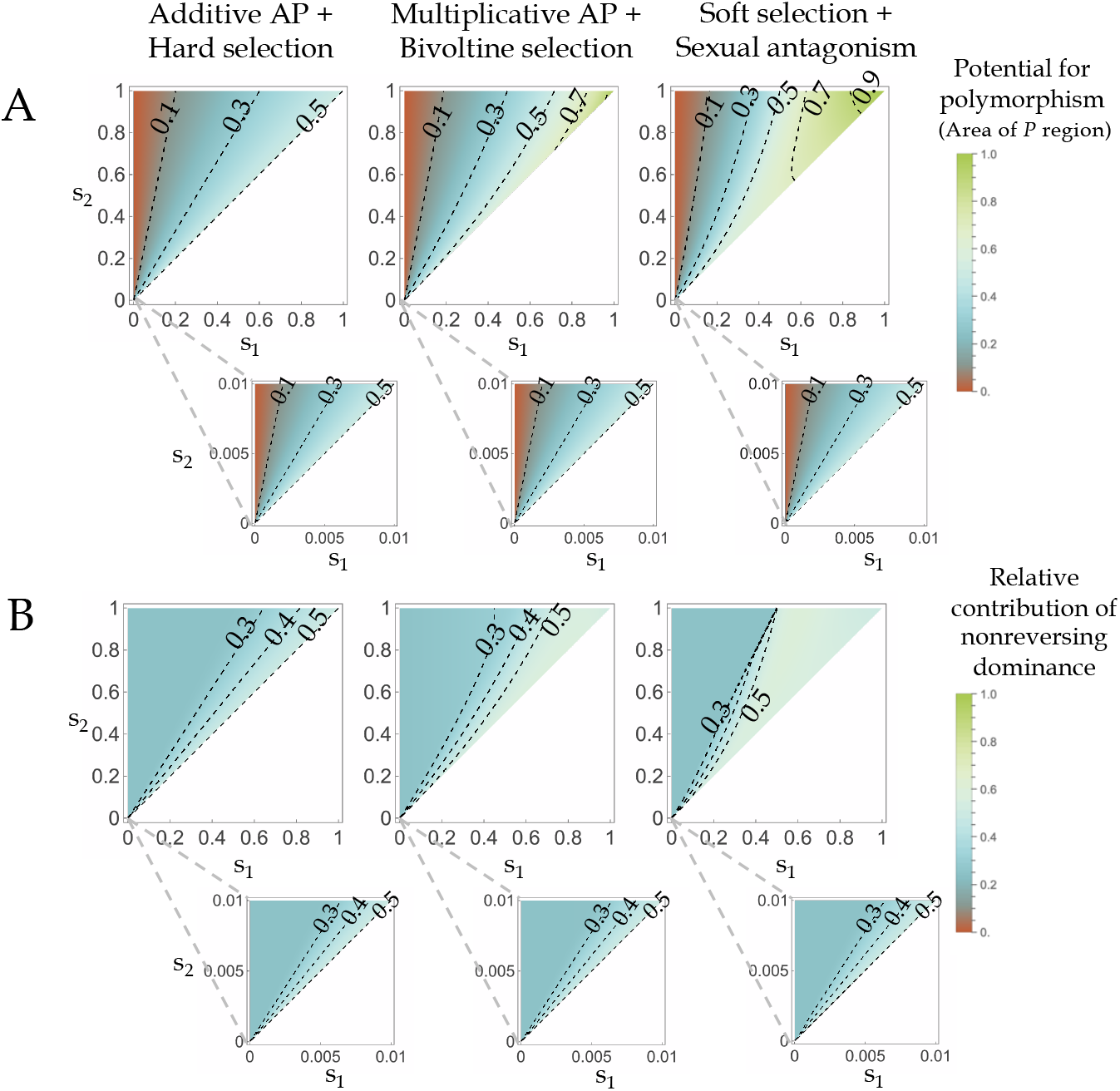
The potential for biallelic polymorphism in six models of antagonistic selection. (A) For each pair of possible selection coefficients (*s*_1_, *s*_2_), contour plots show the size of the region *P* that confers protected polymorphism on the unit square of dominance coefficients. Recall the assumptions *s*_1_ ≤ *s*_2_ (blank areas) and 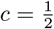 under multiple-niche selection. (Top row) General selection coefficients. (Bottom row) “Weak” selection coefficients, ranging from 0 to 0.01. (B) The relative contribution of non-reversing allelic dominance to the overall potential for polymorphism. For each pair of possible selection coefficients (*s*_1_, *s*_2_), contour plots show the combined size of the subregions formed from the intersection of *P* with the two quadrants representing A-allele dominance and a-allele dominance (relative to the size of the entire region *P*). Recall the assumptions *s*_1_ ≤ *s*_2_ (blank areas) and 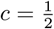 under multiple-niche selection. (Top row) General selection coefficients.(Bottom row) “Weak” selection coefficients, ranging from 0 to 0.01.

The relative contribution of non-reversing dominance to the overall potential for polymorphism is given by the relative area of those subregions of *P* lying in the quadrants that represent a constant direction of dominance for the *A*-or *a*-alleles. This contribution is invariably at least 25%, and grows to 50% with increasing symmetry of selection coefficients (Fig. 2B). Contributions greater than 50% from non-reversing dominance are possible for nearly symmetric (*s*_1_, *s*_2_) regimes under strong selection (with the exception of the Additive AP and Hard selection models).

### Heterozygosity and allele frequency differentiation

The range of equilibrium *a*-allele frequencies 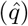 is typically substantial in all models, with the main determinant being the asymmetry in the selection coefficient pairs (Fig. 3, Fig. S2). Two-fold asymmetries not only halve the size of the *P* region, they also push the *a*-allele frequency to values of ∼ 70–90% for much of the space of stabilizing dominance schemes. With schemes that are less than two-fold asymmetric, the space of dominance schemes conferring high equilibrium heterozygosity is seen to be rather broad, with ample contributions from reversing and non-reversing quadrants.

**Fig 3.**
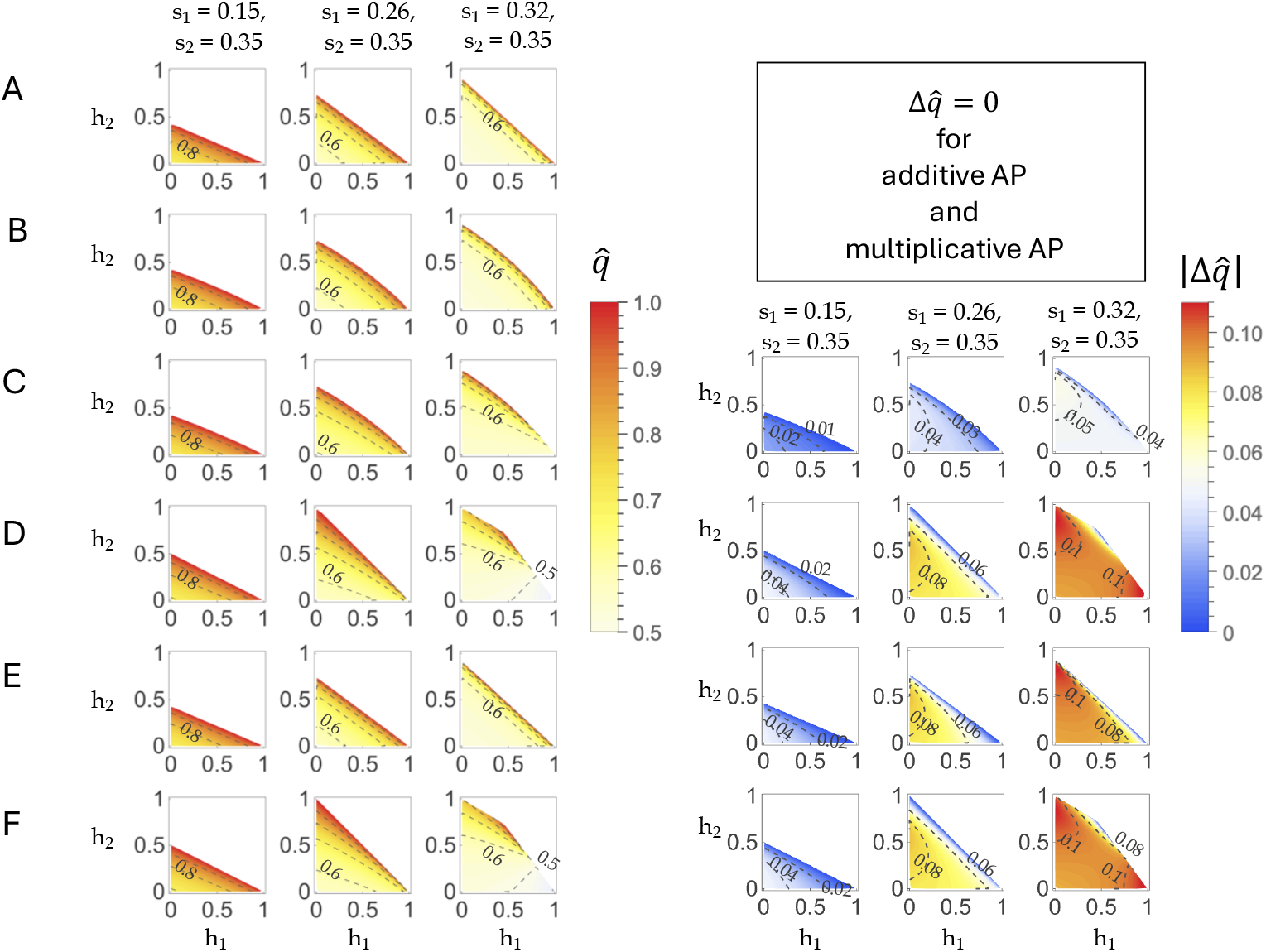
Properties of single-locus equilibria subject to strong antagonistic selection. (Left) Contours (in increments of 0.1) plot the exact equilibrium frequency of the *a*-allele 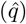 among the total population of zygotes at the start of the generation (or the start of the year under bivoltinism), given the set of strong selection coefficients listed on the column label. Parameter sets (*n* = 2500) were obtained by uniform random sampling from the region *P*. (A) Additive AP, (B) Multiplicative AP, (C) Bivoltine selection, (D) Soft selection 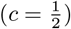. (E) Hard selection 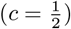, (F) Sexual antagonism. (Right) Contours (labeled) plot the absolute allele-frequency differences 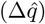 between focal subgroups of the population. (C) Season 1 vs. season 2 frequencies under bivoltine selection; (D–E) niche 1 vs. niche 2 post-selection frequencies under soft and hard selection 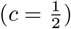 (F) male vs. female post-selection frequencies under sexual antagonism.

Beneficial reversal tends to maximize heterozygosity in all models, with the exception of roughly symmetric pairs of large selection coefficients under soft selection and sexual antagonism (Fig. 3D,F); in these latter models, near-50% *a*-allele frequency coincides with dominance of the *a*-allele. Under weak selection, all six models generally deviate only slightly in their 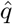 as compared to the additive-AP result (Fig. S3).

Another important property of single-locus equilibria in these models is the absolute allele-frequency differentiation (aAFD, 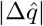) between subgroups of the total population (see Models). Under additive and multiplicative AP, the population does not divide into any relevant subgroups between which 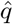 might be differentiated. Under bivoltine selection, there is the difference in 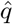 between seasons 1 and 2; under the soft and hard selection models, there is the difference in 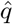 between niches subsequent to zygotic selection; and finally, under sexual antagonism, there is the difference in 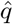 between the sexes.

Greater than two-fold asymmetry of (*s*_1_, *s*_2_) results in beneficial reversal being associated with the maximum aAFD, while more symmetric values result in non-reversing dominance occupying the role of maximizing aAFD. These conclusions hold irrespective of the strength of selection (compare Fig. 3C–F and Fig. S2C–F).

Two additional conclusions regarding the distribution of aAFD across the space of stabilizing dominance schemes merit attention. First, there is a two-fold difference in the magnitude of aAFD between bivoltine selection and the other models. Temporal selection is therefore a milder cause of allele-frequency differentiation as compared to spatial and sexual sources of antagonism. Second, for strong selection with roughly symmetric selection coefficients, both of the non-reversing quadrants (A-allele dominance or *a*-allele dominance) are seen to maximize aAFD under soft selection and sexual antagonism, while in the other models it is dominance of the less-fit A-allele that is the sole maximizer of aAFD (Fig. 3). This pattern does not completely hold under weak selection (again assuming roughly symmetric *s*_1_, *s*_2_), as dominance of the less-fit allele is uniformly associated with maximizing aAFD for all models (Fig. S2).

### Quantitative genetics of fitness at multilocus linkage equilibrium

Complete beneficial reversal is not always associated with the greatest amount of additive genetic variance for fitness characters at an equilibrium locus (Fig. 4A, Fig. S4). Roughly symmetric selection coefficients result in non-reversing dominance acting to maximize 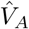 by a factor of up to 1.5× compared to the level of 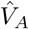 maintained by complete reversal; indeed, approximately half of the *P* region is associated with levels of 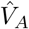 that exceed the complete reversal case. As the asymmetry of selection coefficients increases, the portion of the *P* region with greater 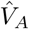 than complete reversal shrinks and practically disappears with greater than two-fold asymmetry in *s*_1_, *s*_2_ (Fig. 4A). The levels of dominance genetic variance in components 1 and 2 never exceed those for the corresponding case of complete reversal, and therefore the total amount of genetic variance in fitness components (summing the additive and dominance genetic variances) that exceeds complete reversal is confined to a smaller region than when considering additive genetic variances alone.

**Fig 4.**
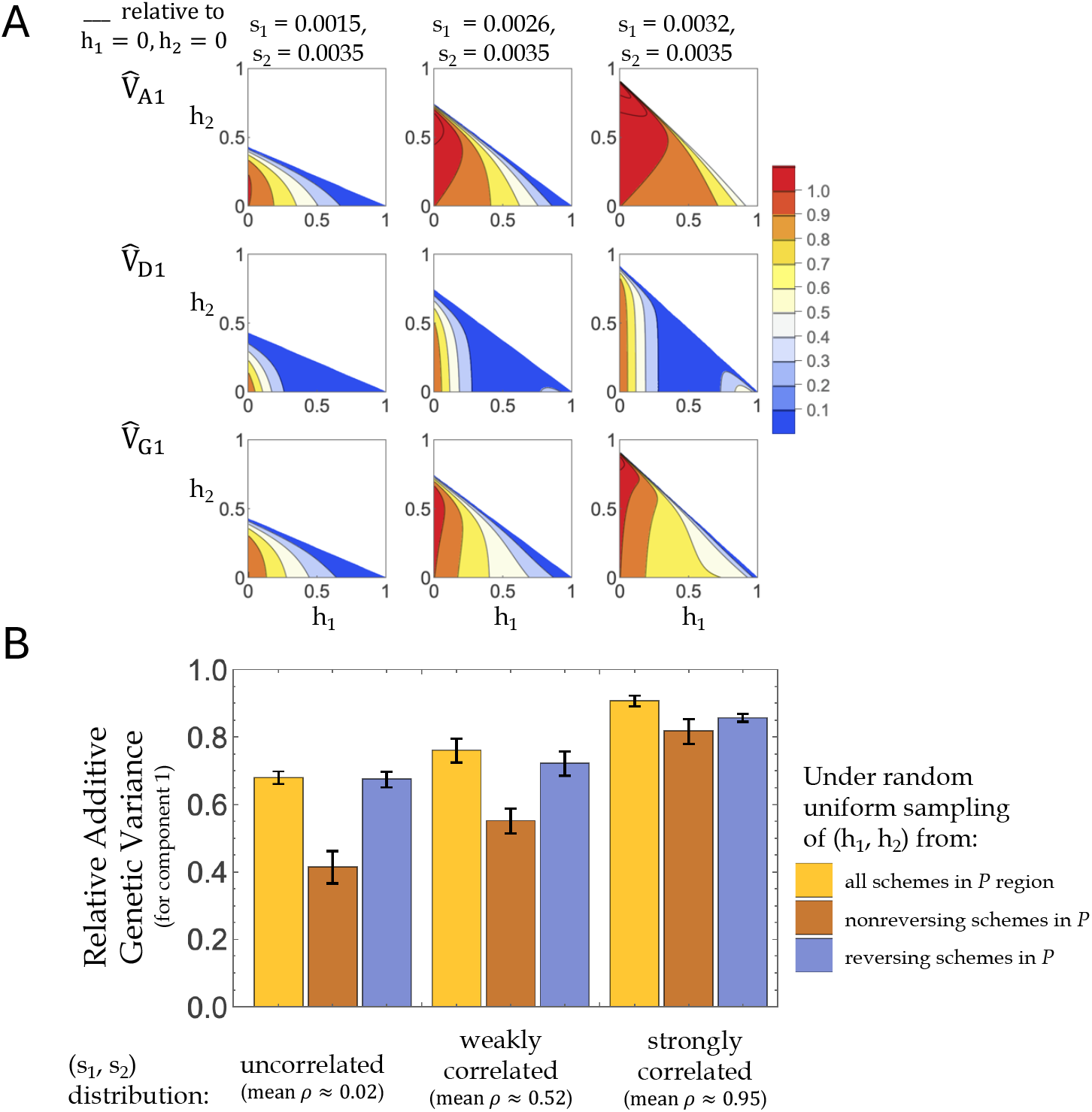
Quantitative genetic analysis of two-component antagonisms at equilibrium. (A) Properties of the single-locus equilibrium under weak selection. Contours plot equilibrium variance terms across the region *P* (for additive AP) given the (*s*_1_, *s*_2_) values labeled on the columns; these terms represent variances relative to the corresponding case of complete beneficial reversal. Regions in red indicate portions of *P* with relative variances greater than those obtained under *h*_1_ = *h*_2_ = 0. (B) Additive genetic variance for component 1 relative to genome-wide complete reversal, 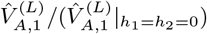, is compared across different sampling schemes for pairs of dominance coefficients (legend), assuming multilocus linkage equilibrium under weak (per-locus) selection. For each of 10 replicates, pairs of selective effects were generated from a bivariate normal distribution (with *ρ* ∈ {0, 0.75, 0.98}) centered at the origin. From these, 250 pairs were randomly chosen from the positive quadrant of the Cartesian plane and multiplied by a factor of 10^−3^, resulting in small selection coefficients (*s*_1_, *s*_2_) at 250 loci (mean correlation coefficients across replicates for the subsamples: *ρ* = {0.02, 0.52, 0.95}). For each (*s*_1_, *s*_2_) pair, a random point from the region *P* (or the subregion indicated by the legend) was drawn with uniform probability; the corresponding relative genetic variance terms were then calculated for each parameter set (*s*_1_, *s*_2_, *h*_1_, *h*_2_) at multilocus linkage equilibrium.

In addition to the single-locus analysis, we investigated the contribution of dominance categories to the multilocus determination of fitness. Assuming that the fitness components are determined by 250 independent loci (i.e. non-epistatic loci with constant linkage equilibrium), the summation of additive variance terms across loci provides the genomewide additive variances 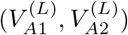 (see Models).

For various levels of correlation between selection coefficients (uncorrelated, *ρ* = 0.02; weak, *ρ* = 0.52; strong, *ρ* = 0.95; where (*s*_1_, *s*_2_) were sampled from the positive quadrant of the bivariate normal distribution centered around the origin), we compared sets of dominance schemes obtained in one of three ways: uniform random sampling of (*h*_1_, *h*_2_) from (i) the entire *P* region; (ii) the non-reversing quadrants of *P*; or (iii) the beneficial reversing quadrant of *P*. These variances were relativized to the optimal case of genome-wide complete beneficial reversal, which maximizes the additive genetic variance in all cases. With uncorrelated selective effects, sampling from the broad *P* region and sampling solely from the beneficial reversing quadrant both result in two-thirds of the standing additive genetic variance in fitness components as compared to the case of genome-wide complete reversal; sampling solely from non-reversing quadrants contributes approximately 40% as compared to the complete reversal reference (Fig. 4B). With stronger levels of correlation between the component-specific selection coefficients, the solely non-reversing samples contribute increasing amounts to the equilibrium additive variance terms so that with 95% correlations between *s*_1_ and *s*_2_, the relative variances are roughly comparable: 0.91 ± 0.02 (uniform in *P*), 0.82 ± 0.04 (non-reversing in *P*) and 0.86 ± 0.01 (reversing in *P*) (mean ratio ± 1 *SD*; Fig. 4B).

## Discussion

Population geneticists have long debated the extent to which allelic variation is maintained by selection in natural populations [36, 37, 38]. The canonical theoretical mechanism is heterozygote advantage [39], in which heterozygous individuals experience a direct gain in fitness relative to homozygotes. Despite its elegance, convincing empirical examples remain relatively scarce [40], and the emphasis on plausible candidate mechanisms has shifted to other models, including antagonistic pleiotropy, temporally and spatially varying selection, sexual antagonism, negative frequency-dependent selection, and selfish genetic elements (e.g. meiotic drive), for which examples are certainly more numerous (see [38]).

A major source of skepticism plaguing several of these mechanisms arises from a common modeling assumption about allelic dominance. Namely, assuming constancy in both magnitude and direction across fitness components imposes stringent restrictions on the permissible values that the selection coefficients can take. This effect is summarized in the familiar “trumpet”-shaped plot of the protection region on the unit square of selection coefficients (Fig. 5), where the interval allowing polymorphism narrows dramatically as selection weakens.

**Fig 5.**
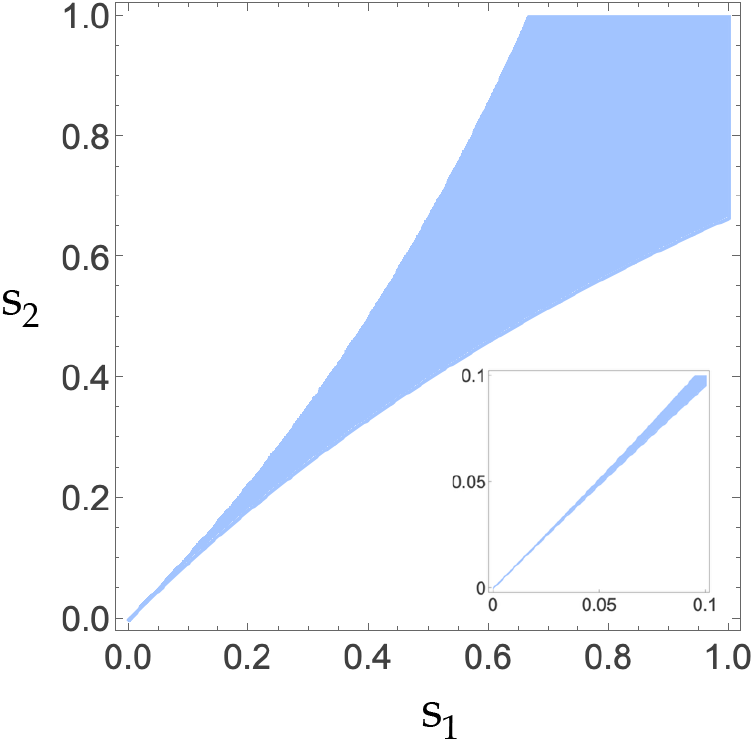
The “trumpet”-shaped region conferring stable polymorphism on the unit square of selection coefficients under additive intralocus effects. Assuming *h*_1_ = *h*_2_ = 0.5 (and general *s*_1_, *s*_2_), the region consistent with protected polymorphism narrows as selection becomes weaker. Even selection coefficients around 10% are characterized by a tight interval (inset).

The problem facing proponents of balancing mechanisms subject to this restrictive effect of the constant dominance assumption is to show that a generalization to component-specific dominance avoids this dramatic shrinkage of the space of permissible parameters. This motivation explains much of the empirical and theoretical attention given to beneficial dominance reversal (theory: [27, 9, 10, 11, 41, 42, 12, 21, 22]; empirical: [43] (copepod *Eurytemora affinis*); [44] (Atlantic salmon, *Salmo salar*); [45] (seed beetle *Callosobruchus maculatus*); [46] (rainbow trout *Oncorhynchus mykiss*); [47] (seaweed fly *Coelopa frigida*); [48, 49, 50, 51, 52, 53] (*Drosophila melanogaster*); reviews: [54, 55, 29]; see Table 1 in [29] for additional empirical cases). Beneficial reversal is undoubtedly the most favorable of the four possible quadrants; in the case of complete reversal (*h*_1_ = *h*_2_ = 0) heterozygotes are entirely free of deleterious effects. However, our results show that the contemporary focus on dominance reversal as a stabilizing mechanism is disproportionate relative to the total space of dominance schemes that maintain polymorphism.

Non-reversing dominance is likely a more common genetic basis for newly arisen antagonistic alleles than reversal. Achieving reversal requires regulatory mechanisms that change the direction of dominance across fitness components (for example through alternating monoallelic expression), whereas non-reversing dominance can arise through subtler shifts in the magnitude of dominance coefficients. Such differences may emerge from, e.g., variation in gene-expression patterns across tissues, sexes, or environments, so that allelic interactions at a locus differ among developmental contexts [53].

These considerations suggest a plausible evolutionary sequence. Antagonistic polymorphisms may initially arise under non-reversing dominance, which our results show already provides substantial potential for stable polymorphism. Subsequent modifications arising at either cis-acting or trans-acting sites may then reduce the fitness costs experienced by heterozygotes by increasing the expression of alleles in contexts where they are beneficial (as argued by Spencer and Priest [56] for sexual antagonism, and Brud [22] for bivoltine selection). Under this view, complete beneficial reversal is likely a derived state, and so the age of allelic variants subject to selective antagonism is predicted to negatively correlate with the distance between their observed dominance schemes and the optimal scheme of maximal reversal (but see [51], who argue that loss-of-function mutants can bring about complete reversal, and so this may form an important class of exceptions to the predicted relationship between allelic age and the intensity of reversal).

Besides reinvigorating the importance of non-reversing dominance, the single-locus results also shed considerable light on the closely similar nature of the various models of selective antagonism. Not only are the sizes of the protection regions exceedingly similar (Fig. 2A; ∼ 80% of dominance schemes having a 10% or greater potential for polymorphism), so are the dominance contributions (Fig. 2B; ∼ 25–50% owing to non-reversals) and the equilibrium allele frequencies (Fig. 3). The models differ primarily in their levels of allele-frequency differentiation (AFD; Fig. 3). Additive and multiplicative AP models lack any basis for AFD, while the two-niche and two-sex models have comparable AFD levels, all roughly two-fold greater than the AFD observed under bivoltine selection. Importantly, under weak selection all six models effectively converge to the additive AP case for all of their properties. Contrary to previous findings that weak selection is unconducive to stabilizing polymorphism owing to AP (see [28]), the differences between weak and strong selection are mainly seen in AFD. The potentials for polymorphism, dominance contributions, and equilibrium heterozygosities are all rather insensitive to the intensity of fitness variation and mainly reflect the asymmetry of selection coefficients rather than their magnitudes. Greater asymmetry in *s*_1_, *s*_2_ results in a decrease of the polymorphic potential and equilibrium heterozygosity, and furthermore two-fold or greater selective asymmetries reduce the relative contribution of non-reversing dominance (to *P*) to nearly its minimum value of 25% (Appendix C).

The convergence of all six models under weak selection also allows a unified treatment of their quantitative genetic consequences. Most importantly, the equilibrium levels of additive genetic variance (the major determinant of a population’s response to selection, together with the phenotypic variance and the “selection differential” that equals the difference in phenotypic means between parents and the total population) are seen to differ depending on (i) the genome-wide distribution of dominance schemes and the strength of correlation in selective effect sizes (Fig. 5B). Interestingly, if antagonistic polymorphisms commonly originate under non-reversing dominance and subsequently evolve toward greater reversal, as suggested above, then additive genetic variance in fitness components should increase over time. Sizeable amounts of additive genetic variance are maintained regardless of which dominance quadrants underlie allelic variation, helping to explain the large amounts of variance in fitness components observed in antagonistic systems [14, 45].

Our results clarify the relationship between antagonistic selection and heterozygote advantage (sensu lato). In Appendix D we show that protected polymorphism under soft selection represents a nonstandard form of heterozygote advantage. While this protection region has been labeled “harmonic mean overdominance,” this terminology can be misleading because the apparent overdominance arises only after heterozygous fitness components are relativized with respect to homozygous components before averaging. This distinction becomes especially important for sexual antagonism. Balanced polymorphism under sexual antagonism has often been treated as conceptually distinct from heterozygote advantage (see Appendix E), but the protection conditions for it (equivalent to the special case of soft selection with *c* = 1*/*2) imply a form of fitness overdominance.

Our focus on two-component antagonism, the simplest case of conflict between fitness components, allows for a thorough dissection of how allelic dominance stabilizes variation. In this setting, dominance relations fall into four broad dominance categories (the four quadrants of the dominance unit square, Fig. 1). Extending the model to include more than two components is possible in principle but comes at the cost of an exponential increase in the number of possible dominance categories. With three components, for example, dominance coefficients occupy a cubic space with eight possible octants; in general the number of possible dominance categories grows as 2^*N*^ for *N* components.

Several other extensions are also possible. The present analysis assumes independence among loci, but epistatic interactions could alter the properties of multilocus equilibria (see [12]). Additionally, our multilocus framework assumes that alleles have consistent directional effects on fitness components. In reality, alleles influence fitness indirectly through their effects on phenotypic traits, and these effects may overshoot or undershoot phenotypic optima. Under these circumstances, allelic effects are rarely universally beneficial or deleterious for a given component, because their fitness consequences depend on the population’s position relative to the phenotypic optimum (see the simulations of [57] for an example of this more complex scenario; cf. [12]). Our results approximate the situation when the phenotypic optima for two traits are sufficiently far from phenotypic means so that overshooting does not occur.

## Conclusion

Our results show that allelic dominance acts in a broadly permissive manner to promote both stable polymorphism as well as the maintenance of additive fitness-component variance in a number of diverse selective scenarios. Previous doubts about the plausibility of antagonistic pleiotropy as a mechanism maintaining allelic variation largely arose from analyses restricted to special dominance relations, namely constant dominance across fitness components and strict additivity. While the stabilizing role of dominance reversal has often been acknowledged, the equally important contribution of non-reversing dominance has rarely been emphasized (but see [11, 41, 12, 22, 58]).

Curtsinger [10], for example, downplayed their simulation results showing stabilizing configurations with both non-reversing and reversing schemes, arguing that the low amounts of dominance genetic variance of phenotypic characters observed in natural populations make antagonistic pleiotropy unlikely to explain large amounts of selective polymorphism. This reasoning is misleading: the relatively small dominance variance in fitness components does not preclude antagonism from contributing substantially to both stable heterozygosity as well as standing levels of additive genetic variance for these traits.

These results suggest several directions for empirical and experimental work. It is of considerable interest to determine to what extent non-reversals and reversals contribute differently to various classes of naturally segregating antagonistic variation. For example, what proportion of the seasonally oscillating SNPs discovered in temperate *Drosophila melanogaster* [59, 60] fall under each category? Does this distribution match that expected for alleles with presumably larger effects than SNPs, such as large inversions (see [53, 52], who find a mixture of contributions from each category for inversions in *Drosophila melanogaster*)? In addition, estimating the relative mutational input for each of the categories would allow for precise calculations of the probability of polymorphism (incorporating mutation rates and stochastic effects in finite populations).

## Supporting information

Supplementary figures are available for download as a pdf: https://github.com/ebrud2/TwoComp_AP/blob/99a7f0cc0528bb024cc3794bcc76c7c2410501f1/Supplement.pdf

**Figure S1**. Life cycle diagrams for various models of antagonistic selection.

**Figure S2**. Equilibrium properties for various models of weak antagonistic selection.

**Figure S3**. Deviance in equilibrium allele frequency between the additive AP result and the other models for weak selection.

**Figure S4**. Quantitative genetic analysis for fitness component 2.

## Acknowledgments

This work was supported by National Institutes of Health/National Institute of General Medical Sciences grant no. R35GM147107.

## Appendix A: Recursion equations for the six focal models

### Model 1: Additive two-trait AP

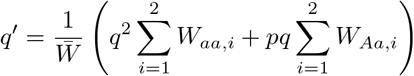

where 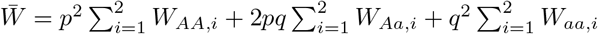.

### Model 2: Multiplicative two-trait AP

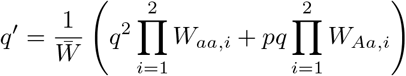

where 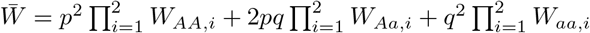.

### Model 3: Seasonal antagonism in a bivoltine population

Following selection in season 1:

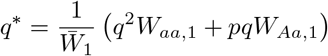

where 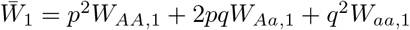.

Following selection in season 2:

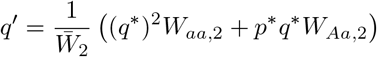

where 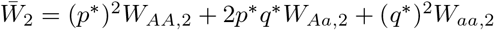.

### Model 4: Antagonistic soft selection between two niches

Post-selection frequencies in niche 1:

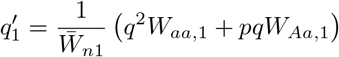

where 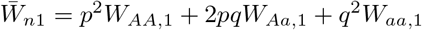.

In niche 2:

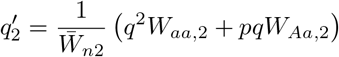

where 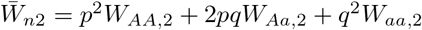.

The allele frequency after mating:

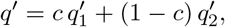

where *c* is the proportional contribution of niche 1 to the common gamete pool.

### Model 5: Antagonistic hard selection between two niches

Using the same within-niche recursions as Model 4, the post-mating frequency is:

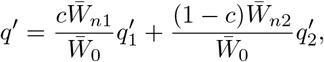

where 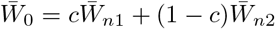.

### Model 6: Sexual antagonism

Following selection in males:

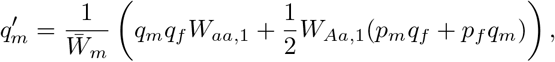

where 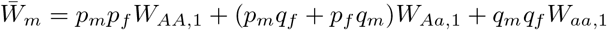.

Following selection in females:

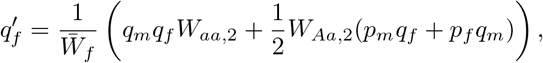

where 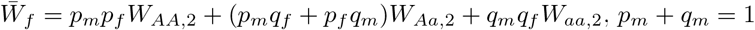, and *p*_*f*_ + *q*_*f*_ = 1. The allele frequency in the total population is:

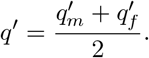

## Appendix B: Absolute areas of the protection region and its dominance category subregions

The formulas below are straightforwardly derived from the geometric picture of the *P* region, as determined by whether *P* intersects 2,3, or 4 quadrants (see figure below):

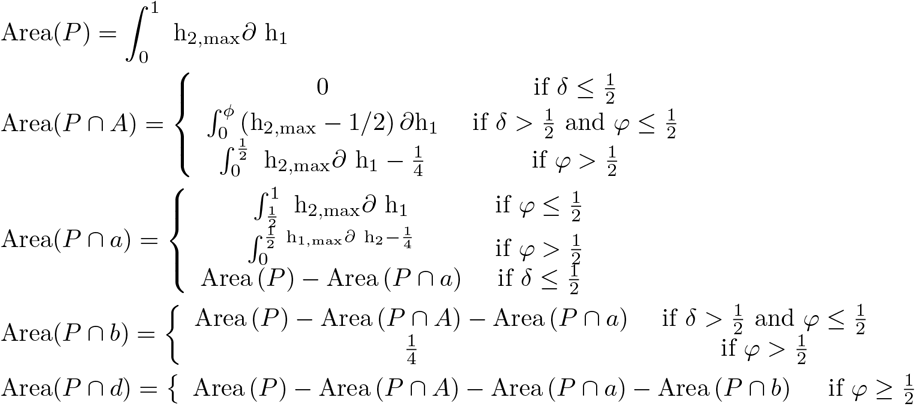

where 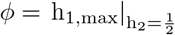 and 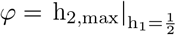. See Table 4 for the explicit areas of the total stabilizing region. Expressions for the intersection areas are not compact expressions but are available in the Mathematica notebook attached with this article.

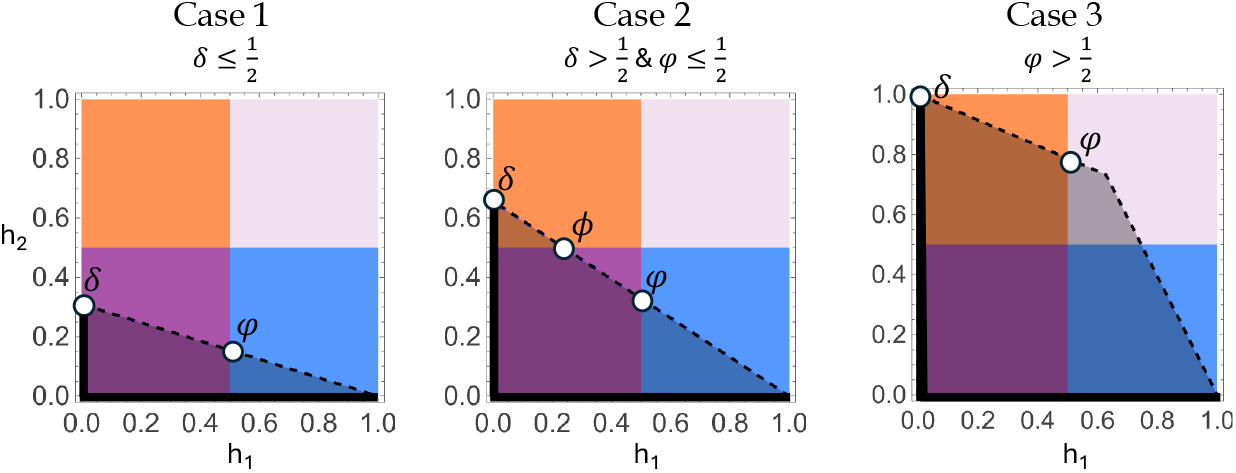

## Appendix C: Weak antagonistic selection reduces to the additive AP model

With small selection coefficients, the single-locus models are unified in that they all approximate the additive AP result.

- The *P* region is given by the right triangle with vertices (0, 0), (0, 1), and (*s*_1_*/s*_2_, 0).
- The upper dominance boundaries are given by:
  - 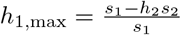
  - 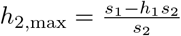
- The size of *P* is straightforwardly derived using trigonometry 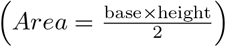 and is equal to 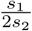.
- The deleterious reversal quadrants makes zero contribution to stability.
- The contributions of the remaining quadrants, relative to the size of *P*, are determined by whether *P* intersects two or three quadrants (Case 1 and Case 2, respectively).

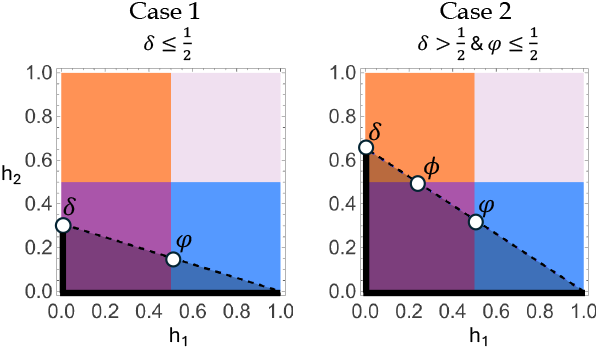

- The points 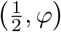 and 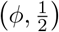 are informative, where
  - 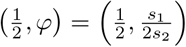 and 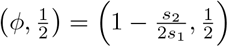
- Case 1: 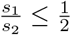
  - a-allele dominance contributes, in absolute area, 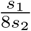 (using the Midpoint Theorem), and with beneficial reversal responsible for the remainder.
  - Notice that this yields 25% as the relative nonreversing contribution.
- Case 2: 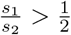.
  - Again, a-allele dominance contributes, in absolute area, 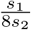.
  - A-allele dominance contributes 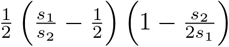 in abs. area.
  - Beneficial reversal contributes the remainder, with a minimum of 50% relative contribution to *P* when *s*_1_ = *s*_2_, rising to 75% for two-fold or greater asymmetry.

## Appendix D: A cautionary note on soft selection and “harmonic mean overdominance.”

The protection region for the soft selection model was originally derived by Levene (1953), who correctly noted that these are “equivalent to the conditions that the weighted harmonic means of the [homozygous niche fitnesses, each relative to the corresponding heterozygous niche fitness – EB & RG] be less than one.” (p. 331; where the weights correspond to the niche size proportions *c*_*i*_). This condition has come to be labeled, somewhat misleadingly, as “harmonic mean overdominance.” To illustrate a confusion that may arise, consider the protection conditions for Additive AP and Multiplicative AP models (the first two rows of Table 2). The Additive AP condition is given as overdominance in the sum of fitness components, or equivalently, overdominance in their arithmetic mean. The Multiplicative AP condition is given as overdominance in the product of fitness components, or equivalently, overdominance in their geometric mean. The Levene (1953) criterion (Table 2; soft selection), however, is neither equivalent to “overdominance in the harmonic mean of fitness components” nor “overdominance in the weighted harmonic mean of fitness components” as one might naively extrapolate from the terminology of the additive and multiplicative cases. This is because the relevant calculation is not an averaging of raw fitness components for each genotype, but rather a calculation focused on the averaging of relativized fitness components. Explicitly:

| Criterion: | Interpretation (for two components): | Correct protection condition? |
| --- | --- | --- |
| Arithmetic mean overdominance | $\frac{\sum_{i=1}^2 W_{AA,i}}{2} < \frac{\sum_{i=1}^2 W_{Aa,i}}{2} > \frac{\sum_{i=1}^2 W_{aa,i}}{2}$ | Yes |
| Geometric mean overdominance | $\sqrt[2]{\prod_{i=1}^2 W_{AA,i}} < \sqrt[2]{\prod_{i=1}^2 W_{Aa,i}} > \sqrt[2]{\prod_{i=1}^2 W_{aa,i}}$ | Yes |
| (weighted) Harmonic mean overdominance | $\frac{1}{\sum_{i=1}^2 \frac{c_i}{W_{AA,i}}} < \frac{1}{\sum_{i=1}^2 \frac{c_i}{W_{Aa,i}}} > \frac{1}{\sum_{i=1}^2 \frac{c_i}{W_{aa,i}}}$ | No! |
| Levene (1953) condition | $\frac{1}{\sum_{i=1}^2 \frac{c_i}{W_{AA,i}/W_{Aa,i}}} > 1$ and $\frac{1}{\sum_{i=1}^2 \frac{c_i}{W_{aa,i}/W_{Aa,i}}} > 1$ | Yes |

Moreover, note that the Levene soft selection condition can justly be characterized in either of two forms, as “overdominance in the weighted arithmetic mean of homozygote-relativized fitness components” (as apparent from Table 2), as well as “overdominance in the weighted harmonic mean of heterozygote-relativized fitness components” (see above). In lieu of the prolixity of these characterizations (and the arbitrariness of highlighting harmonic averaging over arithmetic averaging and vice versa), we propose “soft selection overdominance” as an adequate label and caution against the naïve interpretation of “harmonic mean overdominance.” See Appendix E for a related note on interpretations of sexual antagonism.

## Appendix E: Clarifying the relationship between sexual antagonism and heterozygote advantage

The original derivation of the protected polymorphism conditions for sexual antagonism (SA) from Kidwell and Clegg et al. (1977) is given by:

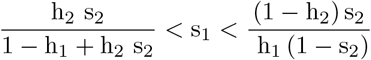

This is equivalent to the statement in Table 2 given in terms of the *W*_*genotype,i*_ parameters of the protection region *R*_*W*_ (i.e. upon translating parameters using Table 1). To our knowledge, the Table 2 version of the SA inequalities is rarely if ever stated. We illustrate below the salubrious effect of this latter formulation on the question of whether balanced polymorphism owing to SA is a form of fitness overdominance (aka heterozygote advantage), which has been denied. Balanced polymorphism at an SA locus has been said to lack a requirement for a “net heterozygote advantage” and, moreover, that beneficial dominance reversal at an SA locus is both (i) what allows for a broadening of the permissible parameter space conferring protection and (ii) constitutes the factor that does impart a net heterozygote advantage (Connallon and Chenoweth 2019, Ruzicka et al. 2025). But this characterization is untenable, since the protection conditions for the SA and soft selection models are identical for equal sized niches, c = 1/2 (Table 2), and moreover, no one disputes that balanced polymorphism under the soft selection model invariably owes to some form of overdominance (we take issue with the label “harmonic mean overdominance” in Appendix D, but end by suggesting alternative formulations that fit squarely under the heading of heterozygote advantage). The overdominant aspect of the soft selection criterion is manifest (esp. in Table 2) as the requirement that heterozygous fitness components have an average relative advantage over the corresponding homozygous components. This applies to SA as an immediate consequence. It is apt, therefore, to refer to balanced polymorphism owing to sexual antagonism as a subcategory of heterozygote advantage models. Furthermore, nonreversing dominance (in *P*) invariably confers a net heterozygote advantage to balanced SA alleles much like reversing dominance (in *P*). (Even deleterious reversal does so for those cases where this scheme is operative, i.e. under strong, nearly symmetric selection). What distinguishes the reversing category is that it contains the point with the greatest fitness boost to heterozygotes, namely complete beneficial reversal, but it is not the case that reversals uniquely convert a disadvantaged situation into one of high fitness for heterozygous genotypes. It is important to note that the above remarks do not imply that the dynamics of sexual antagonism are equivalent to a special case of soft selection (c =1/2). Despite the commonality of the protection region, these models certainly differ. Firstly, the state variables for the gametic frequencies in the SA model are differentiated by sex (*q*_*m*_, *q*_*f*_) and are generally unequal between egg and sperm pools, while gametic frequencies under soft selection are described by the single variable, q (the frequency of the a-allele in the common gamete pool; Figure S1). Secondly, and as a consequence of this prior fact, the distribution of genotypic frequencies under sexually antagonistic selection generally violates Hardy-Weinberg proportions (excess of heterozygotes), while under the soft selection scenario, the niche-specific genotypes among zygotes (i.e. prior to selection) always conform to the Hardy-Weinberg expectation. (See also Connallon et al. 2018 who pointed out the mathematical equivalence between the haploid models of soft selection (c = 1/2) and sexual antagonism).

