## Supplemental Figures for "The geometry of dominance shows broad potential for stable polymorphism under antagonistic pleiotropy"

**Supplement**

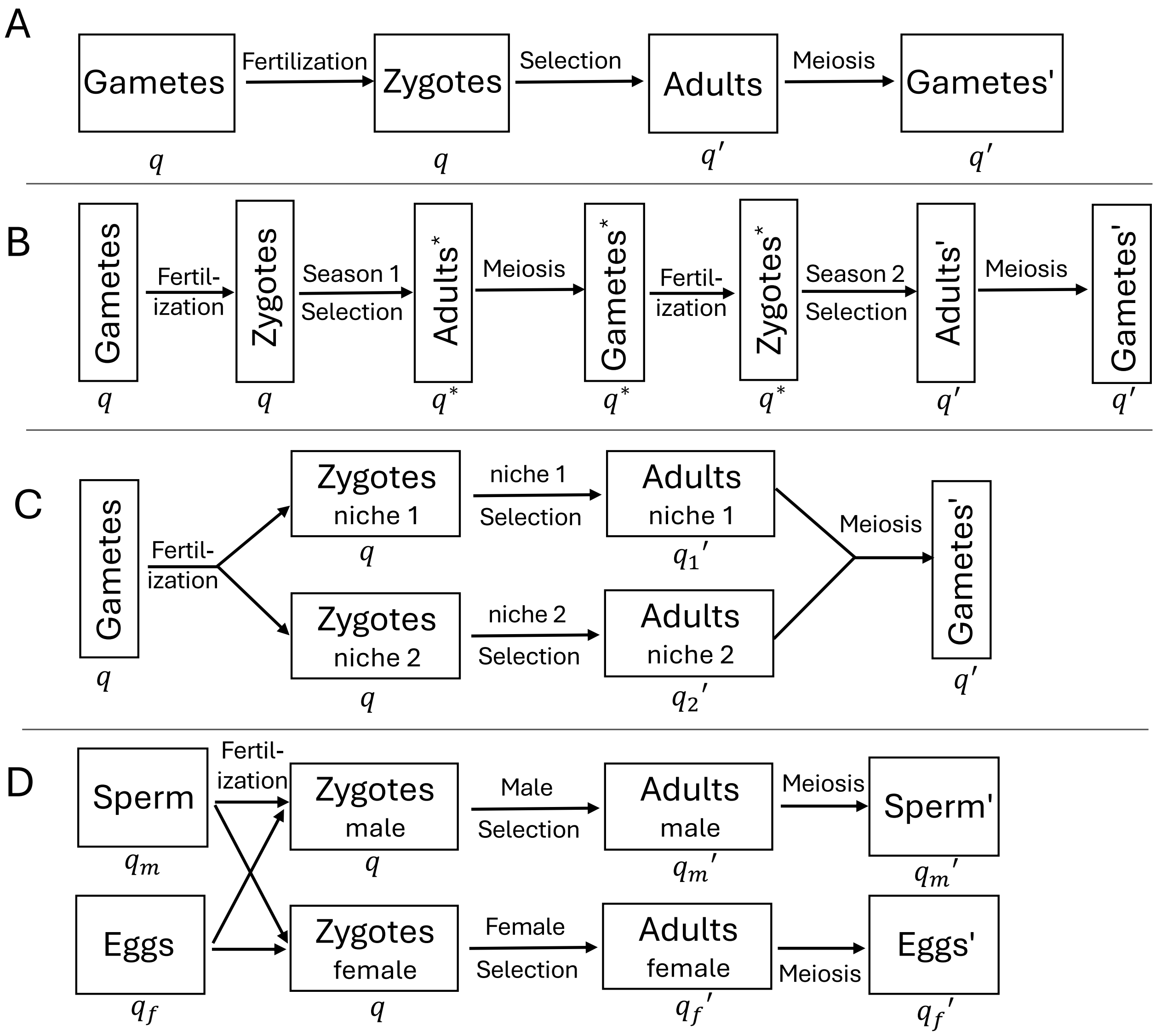


**Figure S1**. **Life cycle diagrams for various models of antagonistic selection.** The frequency of the a-allele is printed below each stage of the life cycle. (A) Additive AP, multiplicative AP. (B) Bivoltine selection. (C) Soft selection, hard selection. (D) Sexual antagonism.

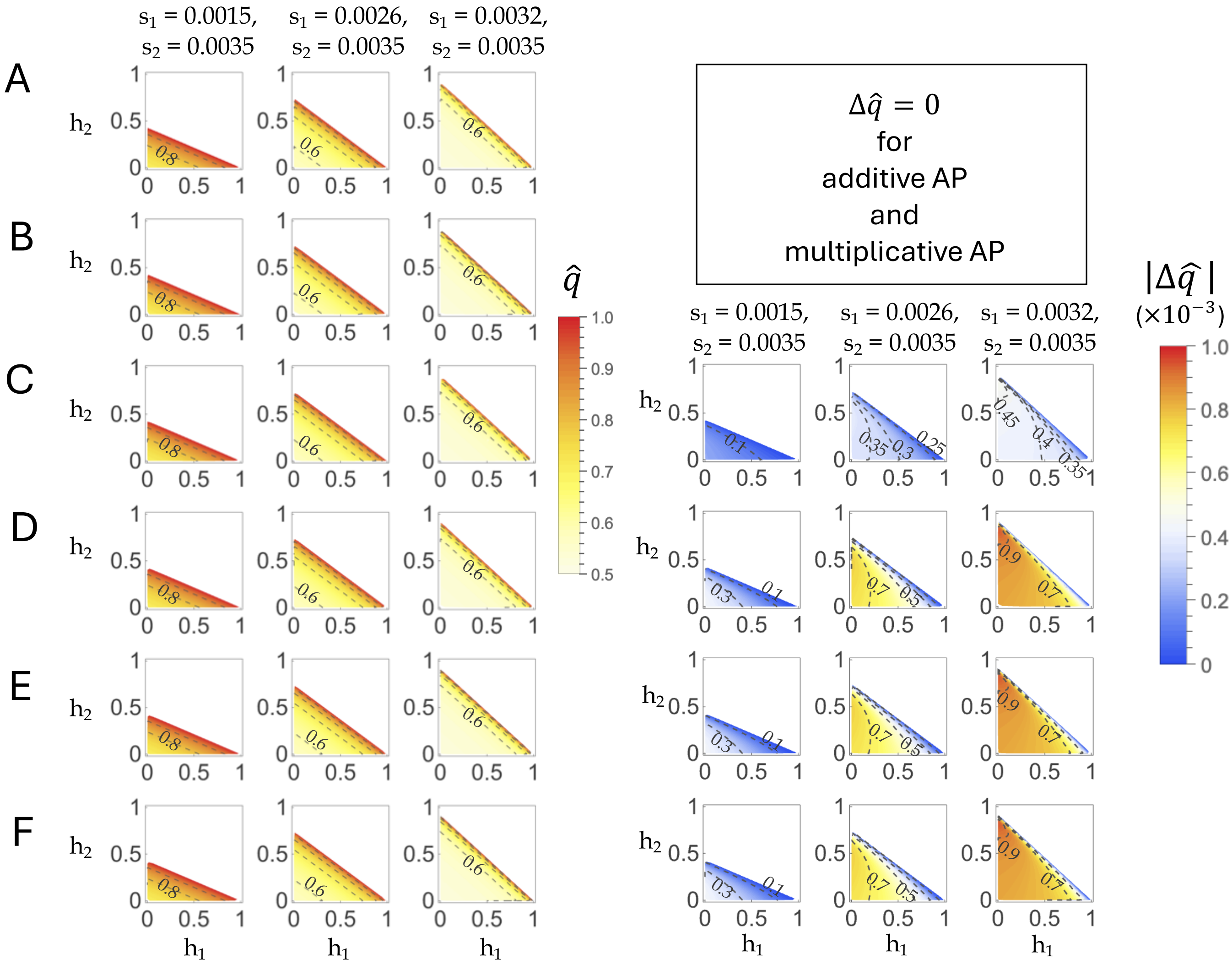


**Figure S2**. **Equilibrium properties for various models of weak antagonistic selection.** (Left) Contours plot the exact equilibrium frequency of the a-allele, $\hat{q}$, among the total population of zygotes at the start of the generation (or the start of the year, for bivoltine selection). (Right) Contours plot the absolute allele frequency differentation between subgroups of the total population. (A) Additive AP, (B) Multiplicative AP, (C) Bivoltine selection, (D) Soft selection (c = 1/2), (E) Hard selection (c = 1/2), (F) Sexual antagonism.

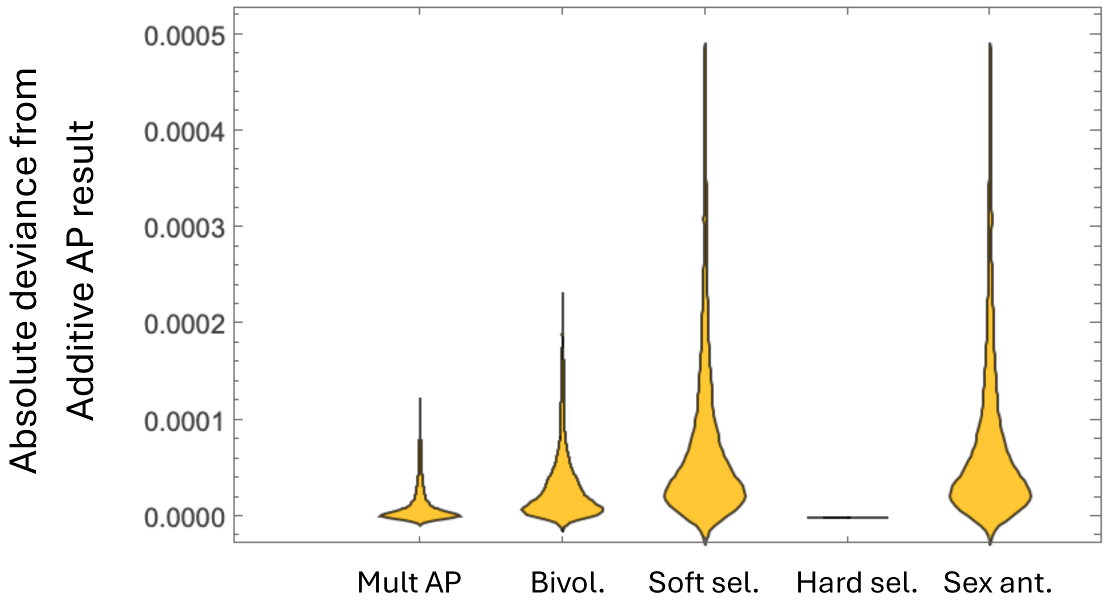


**Figure S3**. **Deviance in equilibrium allele frequency between the additive AP result and the other models for weak selection.** The absolute difference in a-allele frequency at equilibrium ($\hat{q}$) between the additive AP result (used as the reference) and the other five models, where the a-allele is censused during the zygotic stage at the start of the generation (or at the start of season 1, for the case of bivoltine selection). The parameters for $\hat{q}$ were generated by random uniform sampling of 2500 points from the $\mathfrak{R}_{\text{hs}}$ region associated with additive AP and assuming $s_1$ and $s_2$ lie in the interval (0.0001, 0.001), where $s_1 \leq s_2$. For soft and hard selection models, c = 1/2.

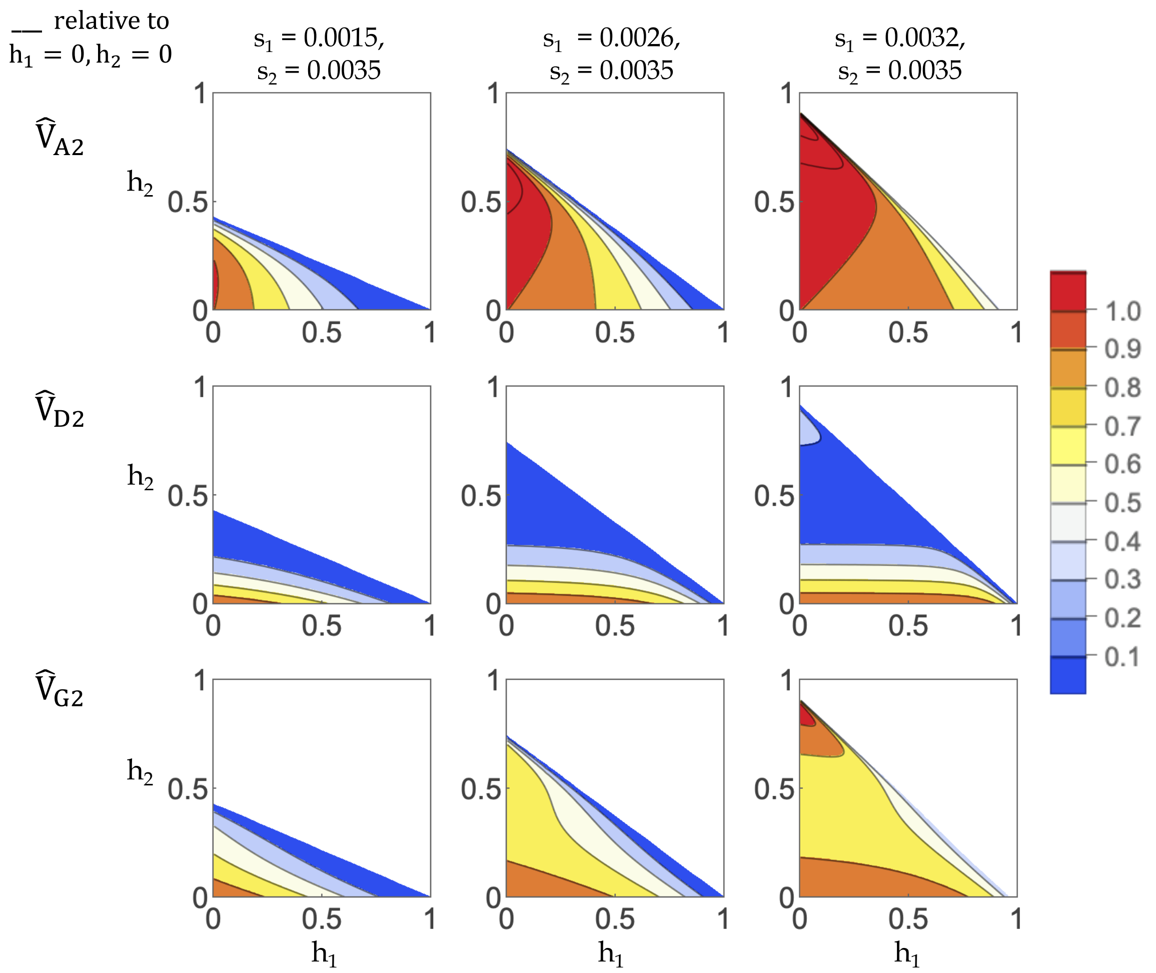


**Figure S4**. **Quantitative genetic analysis for fitness component 2.** See caption for Figure 4A, which had the corresponding plots for fitness component 1.
